# Genome wide association mapping of *Pyrenophora teres* f. *maculata* and *Pyrenophora teres* f. *teres* resistance loci utilizing natural Turkish wild and landrace barley populations

**DOI:** 10.1101/2021.06.07.447398

**Authors:** Shaun J. Clare, Arzu Çelik Oğuz, Karl Effertz, Roshan Sharma Poudel, Deven See, Aziz Karakaya, Robert S. Brueggeman

## Abstract

Unimproved landraces and wild relatives of crops are sources of genetic diversity that were lost post domestication in modern breeding programs. To tap into this rich resource, genome wide association studies in large plant genomes have enabled the rapid genetic characterization of desired traits from natural landrace and wild populations. Wild barley (*Hordeum spontaneum*), the progenitor of domesticated barley (*H. vulgare*), is dispersed across Asia and North Africa, and has co-evolved with the ascomycetous fungal pathogens *Pyrenophora teres* f. *teres* and *P. teres* f. *maculata*, the casual agents of the diseases net form of net blotch and spot form of net blotch, respectively. Thus, these wild and local adapted barley landraces from the region of origin of both the host and pathogen represent a diverse gene pool to identify new sources of resistance, due to millions of years of co-evolution. The barley - *P. teres* pathosystem is governed by complex genetic interactions with dominant, recessive, and incomplete resistances and susceptibilities, with many isolate-specific interactions. Here we provide the first genome wide association study of wild and landrace barley from the Fertile Crescent for resistance to both forms of *P. teres*. A total of 14 loci, four against *P. teres* f. *maculata* and ten against *Pyrenophora teres* f. *teres,* were identified in both wild and landrace populations, showing that both are genetic reservoirs for novel sources of resistance. We also highlight the importance of using multiple algorithms to both identify and validate additional loci.

## INTRODUCTION

Net blotch, caused by the ascomycetous fungal pathogen *Pyrenophora teres* (anamorph: *Drechslera teres*) is an economically important disease of barley worldwide. *P. teres* f. *teres* (*Ptt*) incites the net form of net blotch (NFNB) and *P. teres* f. *maculata* (*Ptm*) incites the spot form of net blotch (SFNB) (Smedegård-Petersen, 1971). Both forms are responsible for large crop losses that typically range between 10-40% when susceptible cultivars are grown, however, under conducive environmental conditions losses can reach 100% (Piening and Kaufmann 1969; Mathre 1982; Moya *et al*. 2018). At least one form of the disease has been reported from all barley growing regions and, in many regions, both forms are present with annual fluctuation in predominance. This presents challenges to breeders, as both *Ptt* and *Ptm* interact with host resistance/susceptibility genes differentially, thus are considered distinct and treated as different diseases when breeding for resistance. However, as further characterization of resistant/susceptibility loci continues, overlaps in host-pathogen genetic interactions in both pathosystems are becoming more prevalent.

Both *Ptt* and *Ptm* occur as genetically distinct populations and can be separated in the field based on lesion morphology. Although these two forms can be hybridized under laboratory conditions (Campbell and Crous 2003), hybridization under field conditions is extremely rare (Campbell *et al*. 2002; McLean *et al*. 2014; Akhavan *et al*. 2015; Çelik Oğuz *et al*. 2018; Poudel *et al*. 2019). However, both forms of *P. teres* undergo form specific sexual as well as asexual reproduction (Karakaya *et al*. 2004; Serenius *et al*. 2005; Akhavan *et al*. 2015; Çelik Oğuz *et al*. 2018, 2019b; Poudel *et al*. 2019). The complex nature of this reproduction system poses serious evolutionary risks for resistance breeding as populations contain diverse effector repertoires that, in different combinations, can rapidly overcome deployed resistances (McDonald and Linde 2002). Yet, the use of resistant barley cultivars is the most environmentally friendly and economically feasible method of NFNB and SFNB control (Robinson and Jalli 1996; Afanasenko *et al*. 2009).

Wild barleys and barley landraces are important sources of resistance against diverse biotic and abiotic stresses (Allard and Bradshaw 1964; Ceccarelli 1996; Yitbarek *et al*. 1998; Ellis *et al*. 2000; Jakob *et al*. 2014; Karakaya *et al*. 2016, 2020). Wild barley (*Hordeum spontaneum*) is known as the progenitor of modern-day barley (*H. vulgare*) and grows naturally in the Fertile Crescent, regions of south and southeastern Turkey, North Africa, and Southwest Asia (Harlan and Zohary 1966; Nevo 1992; Zohary and Hopf 2000). Barley landraces gave rise to modern barley varieties (Thomas *et al*. 1998) as they were subjected to natural and artificial selection for the last 10,000 years (Ceccarelli and Grando 2000). Within the barley center of origin, Turkey is located at the ancestral hub of barley diversification regions including the Mediterranean, Horn of Africa, and the Tibetan Plateau (Muñoz-Amatriaín *et al*. 2014; Poets *et al*. 2015), with diverse barley landraces still used by Turkish farmers (Helbaek 1969; Kün 1996; Pourkheirandish and Komatsuda 2007; Ergün *et al*. 2017). Turkey is also at the center of origin of the *P. teres* pathogens, *Ptt* and *Ptm*. Barley lines can show differential responses to either or both forms of net blotch due to distinct yet complex genetic interactions with each form. Many barley genotypes may be resistant to the majority of isolates of one form yet susceptible to the alternate form of most isolates (Bockelman *et al*. 1983; Grewal *et al*. 2012). Thus, when breeding for resistance, the two forms of net blotch are treated as separate diseases (Liu *et al*. 2011; Usta *et al*. 2014; Yazıcı *et al*. 2015).

In barley, genetic resistance to *P. teres* was first reported in 1928 by Geschele. Since both forms of net blotch were not described at that time, it was assumed that the net form was used. In the first studies to genetically characterize barley – *P. teres* interactions, Tifang was the resistant parent in the Tifang × Atlas cross (Schaller 1955). Mode and Schaller (1958) identified the resistance genes *Pt_1_*, *Pt_2_* and *Pt_3_* segregating in this cross. Bockelman et al. (1977) revised the naming of *Pt_1_*, *Pt_2_* and *Pt_3_* resistance genes and described the *Rpt1* (*Pt_1_* and *Pt_2_*), *Rpt2* (novel) and *Rpt3* (*Pt_3_*) loci on chromosomes 3H, 1H and 2H, respectively. Further studies identified *Rpt4* on chromosome 7H (Williams *et al*. 1999, 2003), *Rpt5* on chromosome 6H, *Rpt6* on chromosome 5H (Manninen *et al*. 2006), *Rpt7* on chromosome 4HL and *Rpt8* on chromosome 4HS (Franckowiak and Platz 2013), however, there are over 340 QTLs previously identified (Clare *et al*. 2020). Manninen et al. (2006) reclassified the locus *Pt_a_* as *Rpt5* on chromosome 6H, and subsequently, three genes/alleles have been characterized at the locus (*Rpt5.f, Spt1.k, Spt1.r*) as dominant resistance or susceptibility genes (Franckowiak and Platz 2013; Richards *et al*. 2016). In multiple barley – *Ptt* genetic interaction studies, it has been shown that the *Rpt5* locus is the most important resistance/susceptibility locus in this system. This complex locus putatively contains multiple resistance as well as susceptibility genes that have been characterized in diverse barley – *P. teres* interactions from around the world (Clare *et al*. 2020). Because the *Rpt5* locus also shows dominant susceptibility in certain barley lines, additional alleles were designated *Susceptibility to P. teres 1* (*Spt1*) by Richards et al., (2016). Further, high resolution genetic mapping and positional cloning efforts have identified *Rpt5* and *Spt1* candidate genes and functional validation is underway (Brueggeman *et al*. 2020).

Resistance to *Ptm* originally appeared to be less complicated when compared to *Ptt* due to the presence of three major loci. These three loci were identified as *Rpt4* on chromosome 7H (Williams *et al*. 1999, 2003), *Rpt6* on chromosome 5HS (Manninen *et al*. 2006) and *Rpt8* on chromosome 4HS (Friesen *et al*. 2006; Franckowiak and Platz 2013). To date, over 140 QTLs have been reported to be implicated in the *Ptm* reaction, which have been collapsed into 36 unique loci, five of which are specific to *Ptm* and the rest showing some degree of overlap with known *Ptt* loci (Clare *et al*. 2020). These five unique loci that are specific to the *Ptm* interaction are *SFNB-3H-78.53* on chromosome 2H (Burlakoti *et al*. 2017)*, QRptm-4H-120-125* on 4H (Tamang *et al*. 2019), *QRptts-5H-106.00* on 5H (Adhikari *et al*. 2019), *QRptm7-*3 on 7H (Wang *et al*. 2015) and *QRptm7-6/QRptm-7H-119-137* on 7H (Wang *et al*. 2015; Tamang *et al*. 2019). Considering that all currently designated resistance/susceptibility loci except for *Rpt2* (only implicated in the *Ptt* interaction) have now been implicated in both *Ptm* and *Ptt* interactions (Clare *et al*. 2020), it is with some caution that it can be concluded that host-pathogen genetic interactions with the two forms and barley should be considered distinct. Thus, for both forms, with the exception of *Rpt6*, numerous researchers have described synonyms of all loci (Clare *et al*. 2020).

Multiple genome wide association mapping studies (GWAS) have investigated NFNB resistance in barley (Wonneberger *et al*. 2017b; Richards *et al*. 2017; Amezrou *et al*. 2018; Adhikari *et al*. 2019, 2020; Rozanova *et al*. 2019; Daba *et al*. 2019; Novakazi *et al*. 2019). A large proportion of the resistance markers associated with NFNB resistance have been localized to the centromeric region of barley chromosome 6H (Richards *et al*. 2017). In GWAS for SFNB resistance, 29 (Wang *et al*. 2015), 27 (Tamang *et al*. 2015), 11 (Burlakoti *et al*. 2017) and one (Daba *et al*. 2019) unique genomic loci were identified. Four important QTLs (*QRptm7-4*, *QRptm7-6*, *QRptm7-7* and *QRptm7-8*) were mapped into a region covering the *Rpt4* locus on chromosome 7HS (Wang *et al*. 2015). Burlakoti et al. (2017) identified a new and important QTL on chromosome 2HS that was predominately found in 6-rowed barley lines as compared to 2-rowed. Daba et al. (2019) also defined a new QTL on chromosome 6H associated with *Ptm* susceptibility in a large number of genotypes, which is a common mechanism in inverse gene-for-gene interactions with this pathogen. Vatter et al. (2017) performed nested association mapping for NFNB and described further interactions at the important *Rpt5* / *Spt1* locus. In barley, the newest approach to identify MTAs with *Ptt* resistance is exome QTL-seq. This approach identified a large number of MTAs on chromosomes 3H and 6H when analyzing resistant and susceptible bulks (Hisano *et al*. 2017). The resistance status of wild barley genotypes and barley landraces to *P. teres* has been reported by several research groups worldwide (Legge et al., 1996; Lakew et al., 1995; Endresen et al., 2011; Silvar et al., 2010; Neupane et al., 2015; Fetch et al., 2003; Jana and Bailey, 1995; Sato and Takeda, 1997; Çelik Oğuz et al., 2017, 2019a, 2019c). However, molecular mapping studies of *P. teres* resistance in wild and landrace barleys have been limited (Yun *et al*. 2005; Vatter *et al*. 2017; Adhikari *et al*. 2019; Gyawali *et al*. 2019, 2020). In this study, four novel loci representing resistance to NFNB were mapped in Turkish wild barley and landraces. This study highlights the importance of surveying wild and unimproved barley lines for sources of resistance that may have been lost during domestication and modern breeding. These studies, focused on diversity in the barley primary germplasm pool, will provide new sources of resistance and associated markers to aid in deploying robust resistances against NFNB and SFNB.

## MATERIALS AND METHODS

### BIOLOGICAL MATERIALS

A total of 295 barley accessions comprised of 193 landraces (*H. vulgare*) and 102 wild barley (*H. spontaneum*) genotypes which were collected from different growing regions of Turkey and maintained at the Field Crops Central Research Institute and Department of Plant Protection, Faculty of Agriculture, Ankara University located in Ankara, Turkey were utilized in the analyses (Çelik Oğuz *et al*. 2017, 2019a). Three virulent *Ptm* isolates (GPS263, 13-179 and 13-167) and three *Ptt* isolates (UHK77, GPS18 and 13-130) collected from different provinces of Turkey (Çelik Oğuz and Karakaya 2017) were used for the phenotypic assessment of the 295 barley accessions.

### PATHOGEN ASSAY PHENOTYPING

Phenotyping of the wild barley (*H. spontaneum*) genotypes and barley landraces (*H. vulgare)* were accomplished according to methods outlined in Çelik Oğuz et al. (2017) and Çelik Oğuz et al. (2019a).

### PCR-GBS LIBRARY PREPERATION AND GENOTYPING BY SEQUENCING

Two independent custom PCR-GBS SNP marker panels containing 365 (Panel 1) (Sharma Poudel *et al*. 2018; Tamang *et al*. 2019) and 1,272 (Panel 2) (Ruff *et al*. 2020) barley SNP markers were used to genotype 295 barley accessions. Marker primers were divided into six and three total primer pools for Panel 1 and 2 for PCR amplification, respectively. PCR amplification and barcoding reactions were performed as described by Sharma Poudel et al. (2018) and Ruff et al. (2020). Briefly, nine primary PCR amplification reactions were performed per sample. Following amplification, equal volumes of primary PCR products were pooled into 96-well plates with each well containing all amplified markers for a given DNA sample. Next, barcoding PCR reactions were performed with a universal barcoding reverse primer and unique forward barcoding primers for each sample. Following barcoding reactions, samples were pooled and purified. A final PCR reaction was performed using sequencing primers to ensure the barcoding reaction was successful. Samples taken before and after the final amplification were run on agarose gel to verify the appropriate product size and amplification. Quantification of the barcoded libraries were performed using the Qubit dsDNA HS assay kit (Life Technologies, Carlsbad, CA). Enrichments were carried out using the Ion OneTouch™ 2 System (Panel 1) and the Ion PI™ Hi-Q Sequencing 200 kit on the Ion Chef (Panel 2). Finally, samples were sequenced on the Ion Torrent Personal Genome Machine™ (Panel 1) and the Ion Proton™ (Panel 2) Systems using two Ion 318™ chips and 3 Ion PI™ chips, respectively, following the manufacturer’s standard protocols. The resulting marker panels were collapsed to eliminate duplicated markers from each panel. The Morex reference map (Beier *et al*. 2017; Mascher *et al*. 2017; Monat *et al*. 2019) and iSelect consensus map (Muñoz-Amatriaín *et al*. 2014) were downloaded from The Triticeae Toolbox (T3) barley database (https://triticeaetoolbox.org/barley/). The Morex reference map was used to determine the absolute marker position, whereas markers not included in the Morex reference map were estimated based on the genetic position of the iSelect consensus map relative to the flanking markers.

### IMPUTATION, FILTERING AND LINKAGE DISEQUILIBRIUM

Due to the heterozygosity present in the natural population, heterozygous calls (5.23%) were included in the analysis. Accessions and markers with more than 30% missing data were removed from analysis, resulting in 282 barley accessions and 530 markers. Missing data was imputed using LinkImpute, which uses a linkage disequilibrium *k-*nearest neighbor imputation (LD-kNNi) method (Money *et al*. 2015) in Trait Analysis by aSSociation, Evolution, and Linkage (TASSEL) 5.2.60 (Bradbury *et al*. 2007). Markers with a minor allele frequency of <0.05 were included in the analysis but were treated with caution based on best practice from the Genomic Association and Prediction Integrated Tool (GAPIT) 3.0 user manual. Linkage disequilibrium was calculated in TASSEL using a window size of 50 markers and an *R^2^* threshold of 0.8 resulting in 522 markers.

### POPULATION STRUCTURE, KINSHIP MATRICES AND MODEL ALGORITHMS

Population structure was accounted for using STRUCTURE analysis and principle component analysis. A total of 522 markers were used for analysis of population structure. The software STRUCTURE v2.3.4 (Pritchard *et al*. 2000) was used to estimate population structure of the barley panel to create a population structure matrix (*Q*) to be used as a covariate. To determine the optimal number of subpopulations, an admixture ancestry model was used with a burnin of 10,000, followed by 25,000 Monte Carlo Markov Chain (MCMC) replications for *k*=1 to *k*=10 with ten iterations. STRUCTURE HARVESTER (Earl and vonHoldt 2012) was used to identify the optimal number of subpopulations using the *Δk* method (Evanno *et al*. 2005). The optimal *k* value was subsequently used to run a new STRUCTURE analysis using a burnin of 100,000 followed by 100,000 MCMC replications. An individual was deemed to be part of a population if the membership probability was >0.8 (Richards *et al*. 2017). Individuals that did not achieve a value of 0.8 were deemed to have admixture ancestry. The final *Q* matrix was used as a fixed covariate in association models. Principle component analysis was conducted in *R* 3.6.3 using GAPIT 3.0 (Wang and Zhang 2018) with default settings. Principle components explaining at least 25% (*PC1*) and 50% (*PC5*) were used for further analysis. A naïve model using only genotypic (Supplemental File 1 and 2) and phenotypic data (Supplemental File 3) and an additional three fixed effect models accounting for population structure (*Q* [Supplemental Figure 4]*, PC1*, and *PC5*) were all performed using the GLM method.

For initial discovery of the most appropriate method to account for random effect in the model, a kinship matrix (*K*) was constructed using the EMMA (Kang *et al*. 2008), Loiselle (Loiselle *et al*. 1995) and VanRaden (VanRaden 2008) algorithms in GAPIT 3.0 (Wang and Zhang 2018) with the MLM model (Yu *et al*. 2006). Based on these results, the EMMA derived *K* matrix (Supplemental File 5) was identified as the most powerful and used for subsequent analysis of mixed models for all isolates that included CMLM (Zhang *et al*. 2010), ECMLM (Li *et al*. 2014) and MLMM (Segura *et al*. 2012). Lastly, SUPER (Wang *et al*. 2014), FarmCPU (Liu *et al*. 2016) and BLINK (Huang *et al*. 2019) algorithms that reconstruct the kinship matrix were used for a total of nine random effect models. Due to the similarity of results, the MLM and MLMM methods were not used in further analysis. Mixed models included combinations to account for population structure (*Q*, *PC1*, and *PC5*), kinship (EMMA *K*) and algorithm methods (CMLM, ECMLM, SUPER, FarmCPU and BLINK) for a total of 15 mixed models per isolate. The mean-squared deviation (MSD) was calculated for each model (Supplemental File 6, Mamidi *et al*. 2011), however visual inspection of QQ plots were done to ensure the model was a good fit. This method was employed as models with the lowest MSD model often had highly correlated observed and expected -log_10_(*p*) values yielding zero significant markers. A Bonferroni correction was calculated at an α level of 0.01 and 0.05 for a -log10(*p*) threshold of 4.72 and 4.02, respectively. Final Manhattan and QQ plots were generated with *R* 3.6.3 package *CMplot 3.5.1* (https://github.com/YinLiLin/R-CMplot).

### QTL IDENTIFICATION

Absolute marker positions were extracted for significant marker trait associations from the Morex reference genome (Beier *et al*. 2017; Mascher *et al*. 2017; Monat *et al*. 2019) and compared to collapsed *P. teres* loci (Clare *et al*. 2020). Significant markers were declared distinct from previously identified loci, i.e. novel, if the nearest neighboring marker that was closer to previously reported loci was not significant or if the gap to currently delimited locus exceeded 10 Mbp in physical distance when no closer marker was present.

## RESULTS

### PHENOTYPIC ANAYLSIS

Wild barley was found to be statistically (Wilcoxon rank sum test) more resistant to *Ptm* isolate 13-179 and *Ptt* isolates GPS18 and UHK77, whereas landrace barley was shown to be statistically more resistant to *Ptm* isolate GPS263 and *Ptt* isolate 13-130 (Figure 1). There was no significant difference between landrace and wild barley to *Ptm* isolate 13-167. Despite UHK77 being statistically different between landrace and wild barley, no significant MTAs were found.

**Figure 1.**
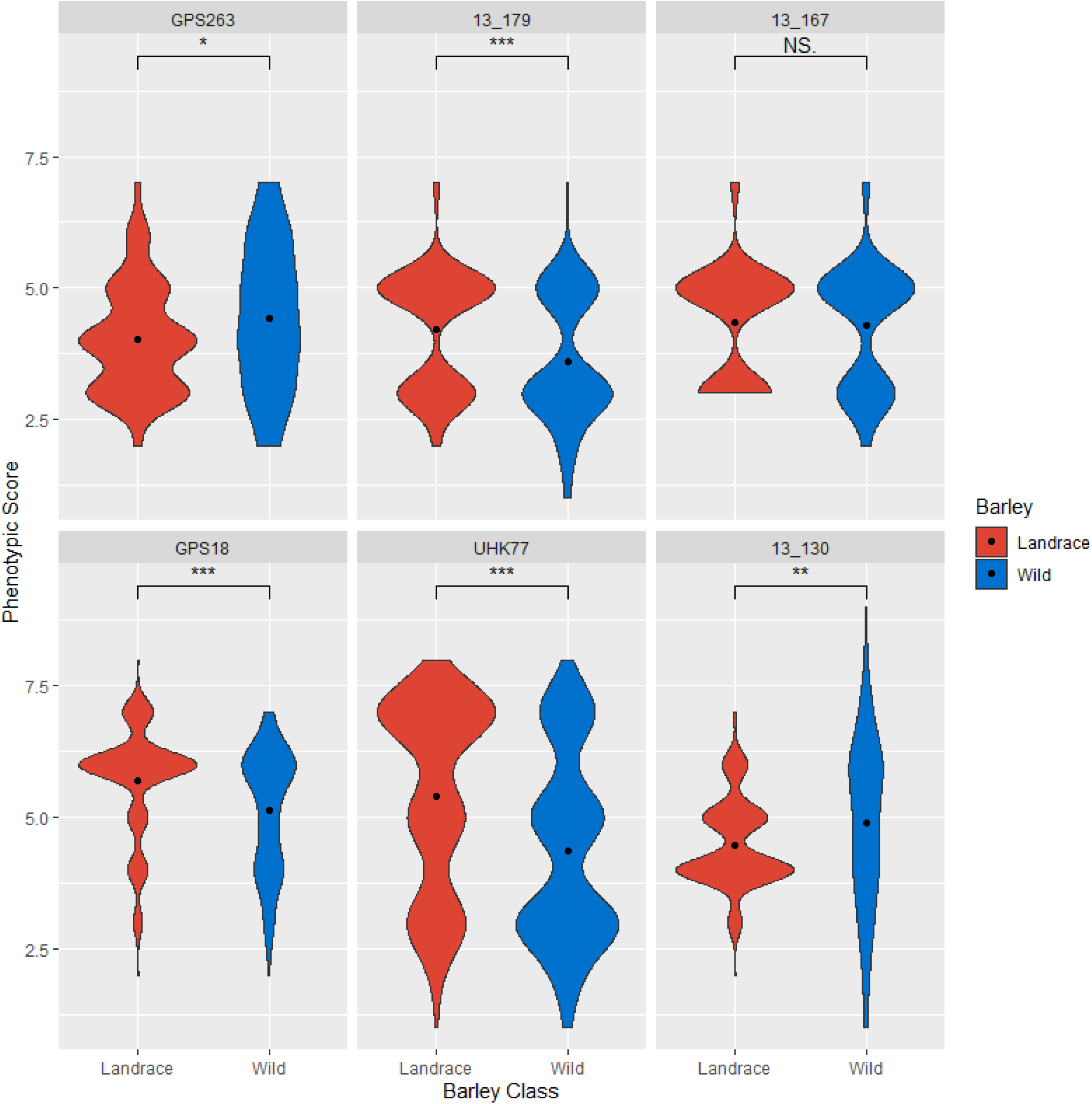
Violin plots of phenotypic distribution of landrace (red) and wild (blue) barley to each *Pyrenophora teres* f. *maculata* (top row) and *P. teres* f. *teres* (bottom row) isolate. Width of the violin indicates the number of accessions with that phenotypic score and the black dot represents the barley class mean. Wilcoxon test significance is indicated by asterisks above each plot.

### MARKER PANEL ANALYSIS

Using a 30% missing data threshold, a total of 282 of the 295 barley accessions phenotyped were adequately genotyped. Using a similarity of individual matrix, all remaining individuals were unique. Similarly, 530 of the 598 collapsed markers were deemed to have sufficient coverage across barley accessions using a 30% missing data threshold and were used in the final analysis of the six *P. teres* isolates. To eliminate markers in linkage disequilibrium, an LD *R^2^* threshold of 0.8 and sliding window of 50 markers was used, resulting in 522 markers for use in the final analysis.

### POPULATION STRUCTURE AND LINKAGE DISEQUILIBRIUM

STRUCTURE analysis identified an optimal *k* value = 2, with 89 and 118 individuals in subpopulations one and two. Subpopulation one consisted of wild accessions, whereas subpopulation two consisted of landraces. A total of 75 barley accessions comprised of both landrace and wild barley had population membership probabilities of less than 0.8 and were deemed to have an admixture ancestry. The first five principle components accounted for 29.36, 9.4, 5.3, 4.0, and 3.7%, respectively, in the principle component analysis. Principle components were selected that accounted for at least 25% (*PC1*) and 50% (*PC5*) when eigenvalues were plotted on a cumulative scale.

### ASSOCAIATION MAPPING ANALYSIS

A total of 24 models were tested on each of the six *P. teres* isolates consisting of three *Ptm* and three *Ptt* isolates. The *Ptm* isolate 13-167 and *Ptt* isolate UHK77 contained no significant markers across all models tested. The *Ptm* isolates GPS263 and 13-179 contained one and three significant markers, respectively. The *Ptt* isolates GPS18 and 13-130 both contained five significant markers.

#### *Ptm* Isolate GPS263

Only one significant MTA was identified with *Ptm* isolate GPS263 on chromosomes 5H based on the second version of the cv. Morex reference genome (Monat *et al*. 2019). The SNP marker 12_20350 located on chromosome 5H at physical position 446449782 was identified at the -log10(*p*) threshold = 4.72 in the K_BLINK_. Marker 12_20350 is embedded within the collapsed *NBP_QRptt5-1* locus (Wonneberger *et al*. 2017a; Clare *et al*. 2020) and was not previously shown to be associated with *Ptm* interactions.

#### *Ptm* Isolate 13-179

The three significant MTAs identified with *Ptm* isolate *13-179* were located on chromosomes 3H, 4H and 5H based on the second version of the Morex reference genome (Monat *et al*. 2019). The marker 11_20866 located on chromosome 3H (physical position 153156749), embedded within the collapsed *QRptms3-2* locus (Wang *et al*. 2015; Wonneberger *et al*. 2017a; Koladia *et al*. 2017; Burlakoti *et al*. 2017; Vatter *et al*. 2017; Rozanova *et al*. 2019; Daba *et al*. 2019; Novakazi *et al*. 2019; Clare *et al*. 2020), was identified at the -log10(*p*) threshold = 4.02 in the K_FarmCPU_ and PC1+K_FarmCPU_ models. The marker 11_10510 is located on chromosome 4H at position 603258307 and was identified at the -log10(*p*) threshold = 4.72 in the K_BLINK_ and K_SUPER_ models and at the -log10(*p*) threshold = 4.02 in the PC5_GLM,_ Q+K_SUPER_, and PC1+K_SUPER_ models. In addition, the 11_10510 marker almost met the significance threshold using the K_FarmCPU_ model. The 11_10510 marker is embedded within the *Rpt8* locus (Friesen *et al*. 2006; Tamang *et al*. 2015; Richards *et al*. 2017; Vatter *et al*. 2017; Daba *et al*. 2019; Clare *et al*. 2020). Lastly, the marker SCRI_RS_160332 is located on chromosome 5H at position 474799503, ∼3.0 Mbp distal to the *Qrptts-5HL.1* locus (Richards *et al*. 2017). SCRI_RS_160332 was identified with the K_FarmCPU_ model at the -log10(*p*) threshold = 4.02.

#### *Ptt* Isolate GPS18

The five significant MTAs identified using *Ptt* isolate GPS18 were distributed across chromosomes 1H, 6H and 7H. The first marker 11_10176 located on chromosome 1H (position 397791042) was identified in the K_FarmCPU_ model at the -log10(*p*) threshold = 4.02. The 11_10176 marker is located 13.6 Mbp distal to the collapsed *NBP_QRptt1-1* (Wonneberger *et al*. 2017a) and 18.2 Mbp proximal to the QTL identified at 57.3-62.8 cM by Rozanova et al. (2019). The marker 11_20754 located on chromosome 1H (position 483805599) was identified in the K_BLINK_ and K_FarmCPU_ models at the -log10(*p*) threshold = 4.72 and 4.02, respectively. The 11_20754 marker is embedded within the *QPt.1H-1* (Vatter *et al*. 2017; Clare *et al*. 2020) locus along with *QRptts-1H-92-93* (Amezrou *et al*. 2018) and a QTL from Tamang et al. (2015). The third marker 12_31282 located on chromosome 7H (position 617741299), is embedded within the collapsed *QTL_UHs_-7H* locus (König *et al*. 2013; Tamang *et al*. 2015, 2019; Wonneberger *et al*. 2017a; Richards *et al*. 2017; Martin *et al*. 2018; Novakazi *et al*. 2019; Clare *et al*. 2020). The 12_31282 MTA was identified using the K_FarmCPU_ model at the -log10(*p*) threshold = 4.72. The fourth marker, 11_20972 on chromosome 6H (position 539551443) is embedded within the collapsed *AL_QRptt6-2* locus (Afanasenko *et al*. 2015; Wonneberger *et al*. 2017b; Vatter *et al*. 2017; Amezrou *et al*. 2018; Clare *et al*. 2020). Marker 11_20972 was identified using the K_BLINK_ model at the -log10(*p*) threshold = 4.02 and also nearly met the significant threshold using the K_FarmCPU_ model. The last significant marker, 12_30545 located on chromosome 7H (54934072) is embedded within the collapsed *QNFNBAPR.Al/S-7Ha* locus (König *et al*. 2013; Tamang *et al*. 2015; Wonneberger *et al*. 2017b; Vatter *et al*. 2017; Amezrou *et al*. 2018; Daba *et al*. 2019; Novakazi *et al*. 2019; Clare *et al*. 2020). Marker 12_20545 was identified at the -log10(*p*) threshold = 4.02 using the K_FarmCPU_ model. Two additional markers, 12_30250 and 12_111942 located on chromosome 3H and 4H, respectively, were almost significant at the -log10(*p*) threshold = 4.02 in the K_FarmCPU_ model. The 12_30250 marker is embedded within the collapsed *QRpts3La* locus (Raman *et al*. 2003; Lehmensiek *et al*. 2007; Liu *et al*. 2015; Tamang *et al*. 2015, 2019; Richards *et al*. 2017; Burlakoti *et al*. 2017; Vatter *et al*. 2017; Daba *et al*. 2019). The 12_11104 marker is located 1.8 Mbp distal from the *Rpt8* locus (Tamang *et al*. 2015; Vatter *et al*. 2017; Clare *et al*. 2020) and 4.4 Mbp proximal to the *QRptm-4H-120-125* locus (Tamang *et al*. 2015, 2019).

#### *Ptt* isolate 13-130

The five significant MTAs identified using *Ptt* isolate 13-130 were distributed across chromosomes 2H, 3H, 6H and 7H. The first marker, 12_11452 located on chromosome 2H (position 34275254) was identified with the K_BLINK_ and K_FarmCPU_ models at the -log10(*p*) threshold = 4.02. The marker 12_11452 has a minor allele frequency less than 5%, however the marker is embedded within the collapsed *SFNB-2H-8-10* locus (Tamang *et al*. 2015, 2019; Wonneberger *et al*. 2017a; Amezrou *et al*. 2018; Adhikari *et al*. 2019; Clare *et al*. 2020). The marker 11_20968, located on chromosome 3H (position 19966889), was identified in the K_FarmCPU_ model at the -log10(*p*) threshold = 4.72. The marker 11_20968 is located 7.6 Mbp distal to the boundary of the collapsed *QPt.3H-1* locus (Vatter *et al*. 2017; Rozanova *et al*. 2019; Daba *et al*. 2019) and 24 Mbp proximal to the boundary of the collapsed *Rpt-3H-4* locus (Tamang *et al*. 2015; Richards *et al*. 2017; Daba *et al*. 2019; Novakazi *et al*. 2019; Clare *et al*. 2020). The marker 12_10662, also located on chromosome 3H (position 553445025), was identified using the K_FarmCPU_ and the Q+K_FarmCPU_ models at the -log10(*p*) threshold = 4.72 and 4.02, respectively. The marker is located 4.6 Mbp distal to boundary of the collapsed *QRpts3La* locus (Liu *et al*. 2015; Tamang *et al*. 2015, 2019; Wonneberger *et al*. 2017a; Richards *et al*. 2017; Burlakoti *et al*. 2017; Vatter *et al*. 2017; Daba *et al*. 2019) and 5.0 Mbp from the boundary of collapsed *Rpt1* locus (Tamang *et al*. 2015; Burlakoti *et al*. 2017; Martin *et al*. 2018; Adhikari *et al*. 2019; Novakazi *et al*. 2019; Clare *et al*. 2020). The last two markers, 11_20714 and 11_11243, were both identified using the K_FarmCPU_ model at the -log10(*p*) threshold = 4.02 and are located on chromosomes 6H (position 489619101) and 7H (position 601974526), respectively. The marker 11_20714 is located 4.1 Mbp proximal to the *QPt.6H-3* locus (Vatter *et al*. 2017) and 11_11243 is embedded within the collapsed *QRptm7-6* locus (Wang *et al*. 2015; Tamang *et al*. 2015, 2019; Wonneberger *et al*. 2017a; Clare *et al*. 2020).

### ENRICHMENT ANALYSIS

Enrichment of either resistance or susceptibility alleles at each MTA were calculated for landraces and wild barley (Figure 2, Supplemental Figure 1). For *Ptm* isolate GPS263 and *Ptt* isolate 13-130, the landraces that were more resistant than the wild barley showed enrichment for the majority of the resistance alleles or a depletion of susceptibility alleles. For *Ptm* isolate 13-179 and *Ptt* isolate GPS18, the opposite was observed as the wild barley showed more resistance and enrichment for the majority of the resistance alleles or depletion of susceptibility alleles.

**Figure 2.**
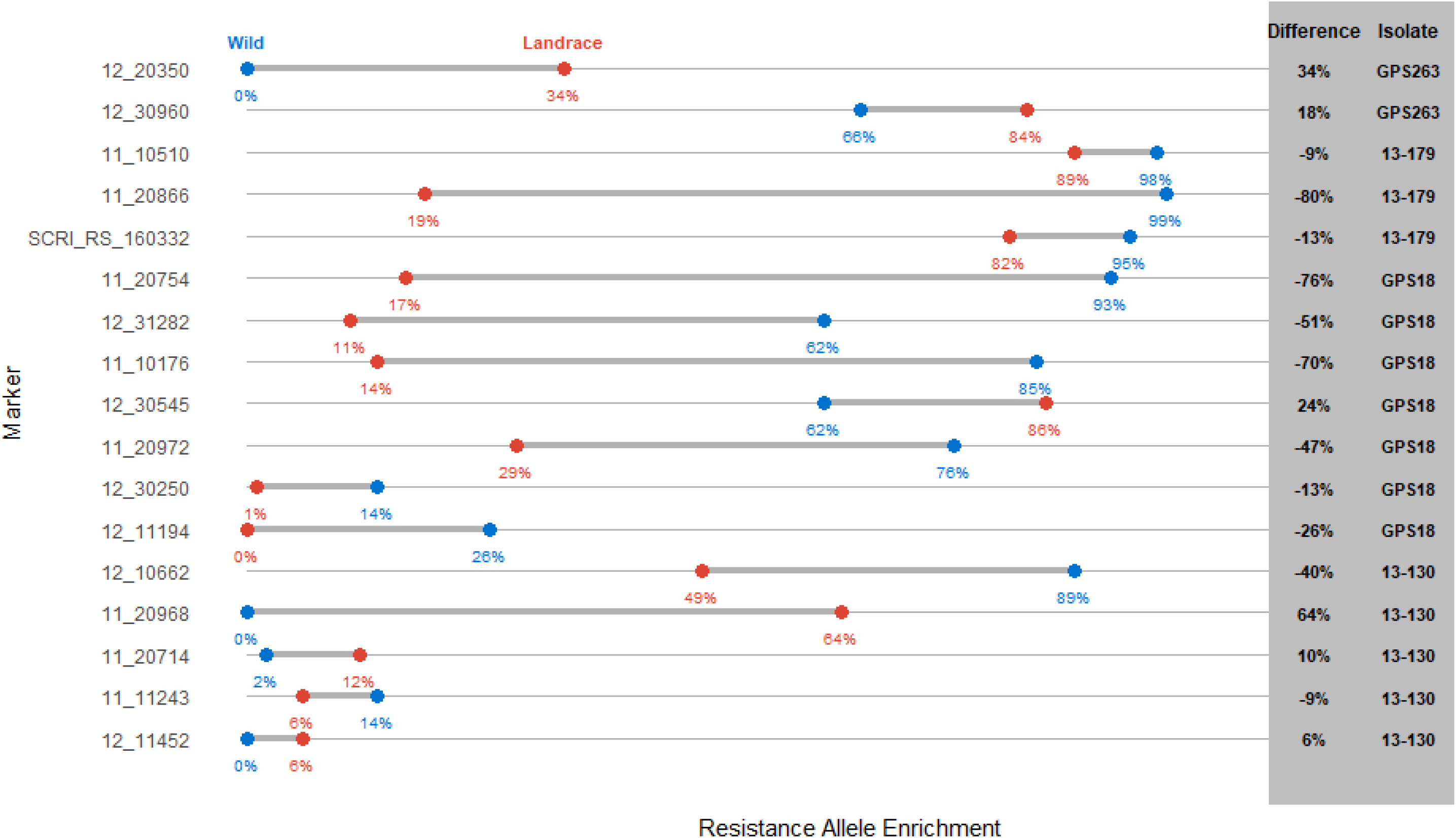
Enrichment dumbbell plot of enrichment of the resistance allele in landrace (red) and wild (blue) barley accession. Percentage is included below each data point and difference between the two barley classes as well as which isolate the loci was identified with.

## DISCUSSION

Both NFNB and SFNB are worldwide threats to barley production and recent evidence shows that both *Ptm* and *Ptt* have evolved to infect and threaten wheat production as well (Tóth *et al*. 2008; Mikhailova *et al*. 2010; Perelló *et al*. 2019). Wild barley and landraces from the origin of cereal domestication represent a rich reservoir of net blotch resistance that could be integrated into elite varieties to help mitigate the threat. Analysis of the phenotypic responses of Turkish wild barley and landraces to regional *Ptm* and *Ptt* isolates showed evidence of landraces under selective pressures by the pathogen during domestication compared with wild barley as seen by the more compact distribution of the phenotypic scores (Table 1, 2, Figure 1). This demonstrates that wild barley harbors additional diversity for net blotch resistance that is not present in the landraces and could be exploited for barley variety development. This is corroborated by enrichment analysis that shows enrichment of the resistant haplotype for marker 11_20866 and 11_20754 in the wild barley lines (Figure 2, Supplemental Figure 1). However, the opposite is true for other loci such as the 11_20968 haplotype near the *QRpt-3H.1* locus, that is located approximately 20 Mbp distal of the domestication gene non-brittle rachis 1, *btr1* on the chromosome 3HS (Komatsuda *et al*. 2004; Wang *et al*. 2019). None of the wild barley accessions analyzed contained the resistance haplotype but it is enriched within landrace accessions (Figure 2, Supplemental Figure 1). The complete lack of the “resistance” marker 11_20968 haplotype in the wild barley could be explained by removal of the “susceptible” haplotype through a selective sweep within close proximity to the *btr1* region during domestication. These results show the importance of surveying both landraces and wild barley accessions since important resistance and/or susceptibility loci that interact in the barley – *P. teres* pathosystem may have been lost or gained through domestication. We have found loci that are unique to wild barley or landraces indicating the importance of analyses of the entire primary barley germplasm pool to identify new sources of resistance for future breeding efforts.

**Table 1.** Phenotypic responses of wild barley and landraces to three *Ptm* isolates with absolute and percentage of accessions in each resistance class. The data is represented as the mean phenotypic score of each barley class to the respective isolate.

| Class | GPS263 |  | 13-179 |  | 13-167 |  |
| --- | --- | --- | --- | --- | --- | --- |
|  | Landrace | Wild | Landrace | Wild | Landrace | Wild |
| Resistant | 6 (3%) | 10 (10%) | 0 (0%) | 3 (3%) | 6 (3%) | 12 (12%) |
| Moderately Resistant | 42 (24%) | 38 (39%) | 65 (37%) | 34 (35%) | 66 (37%) | 50 (51%) |
| Intermediate | 42 (24%) | 28 (29%) | 105 (59%) | 57 (58%) | 100 (56%) | 35 (36%) |
| Moderately Susceptible | 86 (49%) | 21 (21%) | 7 (4%) | 4 (4%) | 5 (3%) | 1 (1%) |
| Susceptible | 1 (1%) | 1 (1%) | 0 (0%) | 0 (0%) | 0 (0%) | 0 (0%) |
| Mean Score | 5.4 | 4.4 | 4.3 | 4.3 | 4.2 | 3.6 |

To date only a handful of studies have utilized wild or landrace barley to map resistance loci using biparental populations (Metcalfe *et al*. 1970; Bockelman *et al*. 1977; Manninen *et al*. 2000, 2006; Williams *et al*. 2003; Koladia *et al*. 2017) or association mapping (Tamang *et al*. 2015; Wonneberger *et al*. 2017a; Richards *et al*. 2017; Vatter *et al*. 2017; Amezrou *et al*. 2018; Adhikari *et al*. 2019; Daba *et al*. 2019; Novakazi *et al*. 2019). Only one study has incorporated both *Ptm* and *Ptt* (Daba *et al*. 2019) and zero have investigated the wild and landrace barley specifically present within the center of origin of the Fertile Crescent. Despite using a reduced marker set (Figure 3) which will reduce the amount of marker trait associations (MTAs) identified (Cui *et al*. 2020), a total of 14 unique MTAs have been identified using modern mapping algorithms (Figure 4). However, the two isolates for which no significant MTAs were identified may be due to the low marker density utilized in these analyses. Four of the MTAs were potentially novel and two that mapped to previously identified *Ptt* resistance loci that had not been reported to be involved in *Ptm* interactions. Additionally, while the remaining MTAs may not be novel, they may represent important alleles that could be incorporated into breeding programs. Thus, the GWAS identified an abundance of net blotch resistance found in this germplasm collected from the barley center of origin.

**Figure 3.**
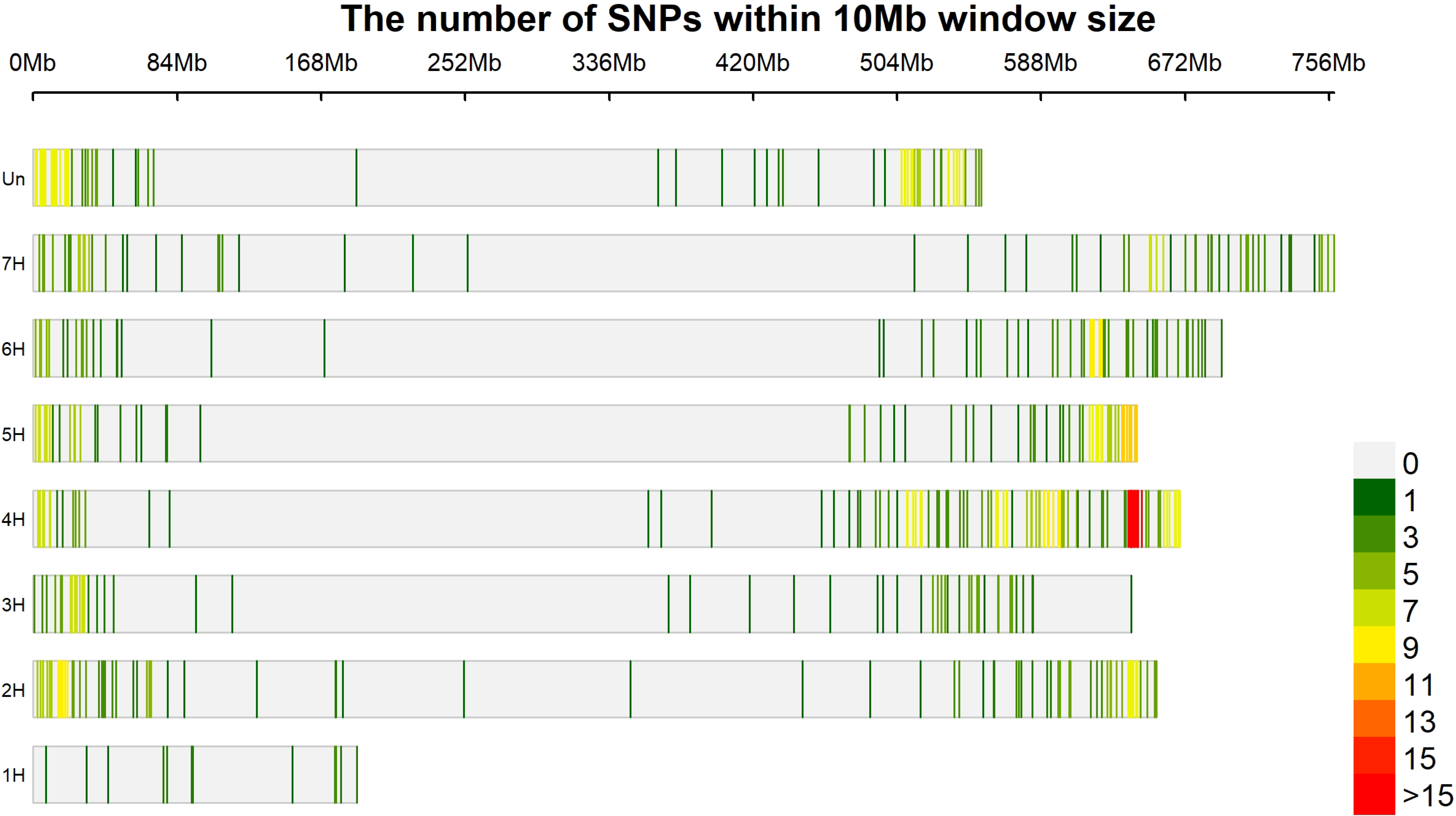
Marker density plot of all markers utilized within this study showing the distribution of markers across the barley genome with a window size of 10 Mb.

**Figure 4.**
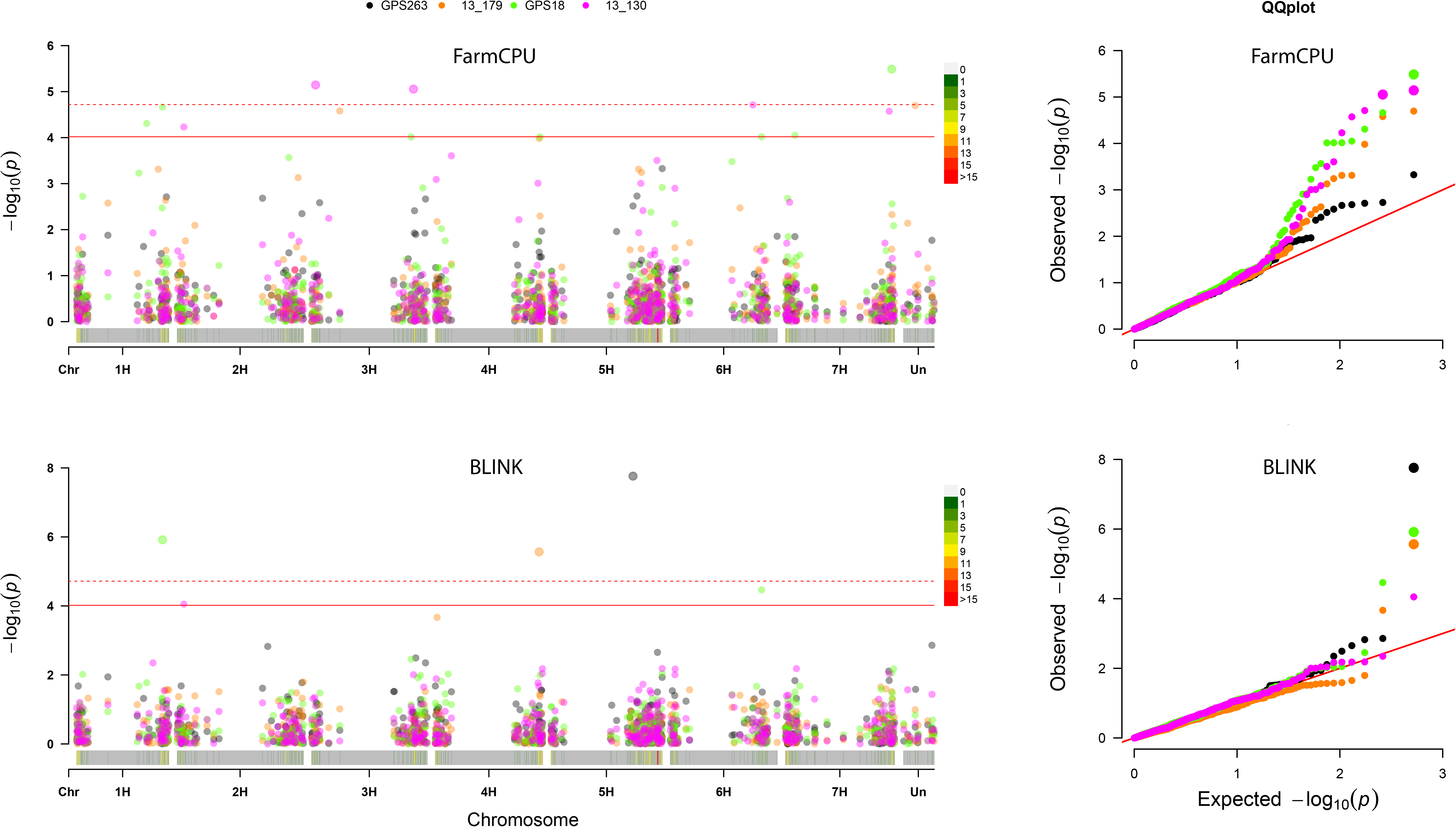
Manhattan and QQ plots for two models that show significant markers for more than one of the *Pyrenophora teres* f. *maculata* isolates GPS263 and 13-179, and *P. teres* f. *teres* isolates GPS18 and 13-130.

In this study barley chromosomes 3H and 7H contained the most MTAs with three, followed by two MTAs on chromosomes 1H, 5H, and 6H and one MTA on chromosomes 2H and 4H (Figure 4, Table 3). Of the 14 MTAs identified, four were identified against the *Ptm* isolates GPS263 and 13-179. The remaining ten MTAs were identified against *Ptt* isolates. When selecting the appropriate GWAS algorithm, the BLINK algorithm identified five MTAs, whereas the FarmCPU algorithm identified eleven MTAs, of which only two MTAs overlapped in both BLINK and FarmCPU algorithms. The SUPER algorithm identified one MTA using *Ptm* isolate 13-179 but this was also identified by the BLINK algorithm. Investigating model selection, the kinship (*K*) model identified 14 MTA, whereas the mixed kinship and population structure (*Q+K*) model only identified one MTA, showing overlap with the *K* model and therefore no unique MTA. Although we would encourage higher marker saturation in future studies to confirm locus novelty, we would suggest continued best practice of testing multiple models (naïve, *K, Q, Q+K*), along with the addition of including all modern association mapping algorithms based on the different set of MTAs identified using BLINK and FarmCPU.

**Table 2.** Phenotypic responses of wild barley and landraces to three *Ptt* isolates with absolute and percentage of accessions in each resistance class. The data is represented as the mean phenotypic score of each barley class to the respective isolate.

| Class | GPS18 |  | UHK77 |  | 13-130 |  |
| --- | --- | --- | --- | --- | --- | --- |
|  | Landrace | Wild | Landrace | Wild | Landrace | Wild |
| Resistant | 1 (1%) | 1 (1%) | 1 (1%) | 7 (7%) | 4 (2%) | 8 (8%) |
| Moderately Resistant | 26 (15%) | 32 (33%) | 103 (58%) | 32 (33%) | 122 (69%) | 44 (45%) |
| Intermediate | 18 (10%) | 14 (14%) | 51 (29%) | 18 (18%) | 35 (20%) | 21 (21%) |
| Moderately Susceptible | 131 (74%) | 51 (52%) | 22 (12%) | 38 (39%) | 16 (9%) | 25 (26%) |
| Susceptible | 1 (1%) | 0 (0%) | 0 (0%) | 3 (3%) | 0 (0%) | 0 (0%) |
| Mean Score | 5.7 | 5.1 | 4.5 | 4.9 | 4.0 | 4.4 |

**Table 3.** Identified significant markers from genome wide association analysis order by isolate identified, followed by chromosome and base pair position. Designation as well as predicted corresponding loci, models used to identify the marker and resistant/susceptibility alleles are also included.

| Loci | Marker | Chr<sup>1</sup> | Position<sup>1</sup> | Allele<sup>2</sup> | Isolate | Corresponding Loci | Models Identified | LOD score<sup>3</sup> |
| --- | --- | --- | --- | --- | --- | --- | --- | --- |
| *QRpt-5H.2* | 12_20350 | 5H | 446449843 | G/A | GPS263 | *NBP_QRptt5-1* | K<sub>BLINK</sub> | 7.76 |
| *QRpt-3H.2* | 11_20866 | 3H | 153156749 | G/A | 13_179 | *QRptms3-2* | K<sub>FarmCPU</sub>, PC1+K<sub>FarmCPU</sub> | 4.58 |
| *QRpt-4H.1* | 11_10510 | 4H | 603258307 | A/G |  | *Rpt8* | K<sub>BLINK</sub>+SUPER, PC5<sub>GLM</sub>,<br>Q+K<sub>SUPER</sub>, PC1+K<sub>SUPER</sub> | 5.57 |
| *QRpt-5H.1* | SCRI_RS_160332 | 5H | 474799503 | A/G |  | *Qrptts-5HL.1* | K<sub>FarmCPU</sub> | 4.70 |
| *QRpt-1H.1* | 11_10176 | 1H | 397791042 | G/C | GPS18 | Novel | K<sub>FarmCPU</sub> | 4.31 |
| *QRpt-1H.2* | 11_20754 | 1H | 483805599 | C/G |  | *QPt.1H-1* | K<sub>BLINK</sub>+FarmCPU | 4.66-5.92 |
| *QRpt-6H.2* | 11_20972 | 6H | 539551443 | T/A |  | *AL_QRptt6-2* | K<sub>BLINK</sub> | 4.46 |
| *QRpt-7H.1* | 12_30545 | 7H | 54934072 | A/G |  | *QNFNBAPR.AL/S-7Ha* | K<sub>FarmCPU</sub> | 4.05 |
| *QRpt-7H.3* | 12_31282 | 7H | 617741299 | C/T |  | *QTL<sub>UHS</sub>-7H* | K<sub>FarmCPU</sub> | 5.49 |
| *QRpt-2H.1* | 12_11452 | 2H | 34275254 | G/A | 13_130 | *SFNB-2H-8-10* | K<sub>BLINK</sub>+FarmCPU | 4.05-4.23 |
| *QRpt-3H.1* | 11_20968 | 3H | 19966889 | A/G |  | Novel | K<sub>FarmCPU</sub> | 5.15 |
| *QRpt-3H.3* | 12_10662 | 3H | 553445025 | A/T |  | Novel | K<sub>FarmCPU</sub>, Q+K<sub>FarmCPU</sub> | 4.22-5.06 |
| *QRpt-6H.1* | 11_20714 | 6H | 489619101 | G/A |  | Novel | K<sub>FarmCPU</sub> | 4.71 |
| *QRpt-7H.2* | 11_11243 | 7H | 601974526 | A/G |  | *QRptm7-6* | K<sub>FarmCPU</sub> | 4.57 |

<sup>1</sup>, Location based on the second version of the Morex assembly (Monat et al., 2019).
<sup>2</sup>, Resistant/susceptible allele
<sup>3</sup>, LOD scores/ranges for BLINK and FarmCPU algorithms only

The 14 MTAs identified in this study were compared to the collapsed loci of Clare et al. (2020) and a similar strategy of determining novel loci was dependent on the significance of the nearest neighbor marker and previous incorporation into a locus. Using this strategy, we have identified four potentially novel loci (*QRpt-1H.1, QRpt-3H.1, QRpt-3H.3, QRpt-6H.1*) in the barley – *Ptt* interaction and two loci previously described only against *Ptt* that had not been previously identified in *Ptm* interactions (*QRpt-5H.1* and *QRpt-5H.2* corresponding to *NBP_QRptt5-1* and *Qrptts-5HL.1,* respectively). The novel loci were detected on chromosomes 1H, 3H and 6H. The *QRpt-1H.1* and *QRpt-6H.1* MTAs were associated in the interaction with *Ptt* isolate GPS18. *QRpt-1H.1* is proximally flanked by the previously reported QTLs *NBP_QRptt1-1* (Wonneberger *et al*. 2017a) and a QTL identified at 57.3-62.8 cM (Rozanova *et al*. 2019) at distances of 13.6 Mbp and 18.2 Mbp, respectively, to the closest boundary of the delimited region of the loci (Clare *et al*. 2020). Markers located 18.2 Mbp proximal and 7.3 Mbp distal to *QRpt-1H.1* were not significant and therefore the MTA was deemed novel. Similarly, the closest locus to *QRpt-6H.1* is *Rpt5/Spt1* (Steffenson *et al*. 1996; Manninen *et al*. 2000, 2006; Raman *et al*. 2003; Read *et al*. 2003; Emebiri *et al*. 2005; Friesen *et al*. 2006; Abu Qamar *et al*. 2008; Grewal *et al*. 2008, 2012; St. Pierre *et al*. 2010; Cakir *et al*. 2011; Gupta *et al*. 2011; O’Boyle *et al*. 2014; Liu *et al*. 2015; Hisano *et al*. 2017; Islamovic *et al*. 2017; Koladia *et al*. 2017; Wonneberger *et al*. 2017b; Richards *et al*. 2017; Vatter *et al*. 2017; Martin *et al*. 2018; Amezrou *et al*. 2018; Adhikari *et al*. 2019, 2020; Rozanova *et al*. 2019; Daba *et al*. 2019; Novakazi *et al*. 2019) located 10.7 Mbp distal, however, a marker 28 Mbp distal to *QRpt-1H.1* and embedded within *Rpt5/Spt1* was not significant, giving us reason to believe the locus is novel. The novel loci *QRpt-3H.1, QRpt-3H.3* and *QRpt-6H.2* were all identified with *Ptt* isolate 13-130. The locus *QRpt-3H.1* is flanked by QTLs located at 12.1-17.4 cM (Rozanova *et al*. 2019) proximal and 53.42 cM (Tamang *et al*. 2019) distal on the Morex POPSEQ map (Mascher *et al*. 2013, 2017), equating to distances of 7.6 Mbp and 24 Mbp. The nearest neighbor markers are 5.2 Mbp proximal and 3.2 Mbp distal and are not significant indicating that this is potentially a novel locus. Similarly the *QRpt-3H.3* locus is flanked by *QRpts3La* (Raman *et al*. 2003; Lehmensiek *et al*. 2007; Liu *et al*. 2015; Richards *et al*. 2017; Vatter *et al*. 2017; Daba *et al*. 2019) and *Rpt1* (Bockelman *et al*. 1977; Graner *et al*. 1996; Richter *et al*. 1998; Raman *et al*. 2003; Cakir *et al*. 2003; Manninen *et al*. 2006; Lehmensiek *et al*. 2007; Martin *et al*. 2018; Adhikari *et al*. 2019; Novakazi *et al*. 2019) approximately 4.6 Mbp proximal and 5 Mbp distal. However, the nearest neighbor markers are not significant at 6.2 Mbp proximal and 1.3 Mbp distal. The last novel locus, *QRpt-6H.2* is 4.1 Mbp proximal to *QPt.6H-3* (Daba *et al*. 2019), however the nearest marker is 8.1 Mbp proximal and not significant.

Two loci that were previously implicated in *Ptt* resistance were also identified as novel MTAs for *Ptm* resistance in this study. The first locus, *QRpt-5H.1,* is located 3.0 Mbp distal to *Qrptts-5HL.1* (Richards *et al*. 2017) with marker SCRI_RS_160332. Additionally, since markers covering the *Qrptts-5HL.1* locus were not included in either panel, and due to the close proximity of *QRpt-5H.1 Qrptts-5HL.1*, we believe that they are the same locus. The second locus, *QRpt-5H.2,* is embedded within the *NBP_QRptt5-1* locus (Wonneberger *et al*. 2017a). The remaining loci are all embedded within previously identified loci (Table 3).

Barley is predominately grown as a feed crop (IBGSC 2012), however in the US where corn and soybeans are subsidized and used as feed crops it has been outcompeted and acreage has significantly dropped. This has pushed feed barley into less than optimal agricultural land due to its adaptability and hardiness (Brueggeman et al., 2020). Quality malting barley demands premium prices because of its use in the multi-billion dollar added value brewing and distilling industry and is now the major class considered in breeding efforts in the US and Europe. However, in some regions of the world where traditional farming practices are still utilized barley is considered an important food crop (Helbaek 1969; Pourkheirandish and Komatsuda 2007; Geçit 2016; Ergün *et al*. 2017). Recent studies predicted that due to the effects of climate change including higher temperatures, altered precipitation patterns, and higher disease pressure (Dawson *et al*. 2015) the world could experience world malt barley shortages to feed the brewing and distilling industry (Xie *et al*. 2018). These predictions are beginning to be seen with the stagnating yields experienced in southern Europe (Dawson *et al*. 2015). Thus, barley breeding must maximize its potential in terms of quality and yield on the land it is currently afforded to sustain the demands for malting. One pillar of support for improving barley would be the introgression of pre-domestication resistance loci that are absent in current breeding programs to prevent substantial losses to net blotch, an important disease effecting barley production across the globe. Here we report on the identification of novel loci from Turkish wild barley and landraces that could be introgressed into elite barley varieties.

## Data Availability Statement

Isolates are available upon request. The authors ensure that all data necessary for confirming the conclusions of the article are present within the article, figures, and tables. Supplemental data has been submitted to Figshare. File SF1 contains genotyping data. File SF2 contains marker positions. File SF3 contains phenotyping data. Files SF4 and SF5 contain population structure and EMMA kindship matrix, respectively. File SF6 contains the mean-squared deviations of each association mapping model tested.

## Author Contributions

ACO, AK and RB conceived the study. ACO and AK carried out phenotyping and DNA extractions. KE and DS carried out sequencing for genotyping data. SC and RSP performed genotyping and analysis. SC wrote the manuscript with contributions from ACO, KE, RSP DS, AK and RB. All authors approved the final manuscript.

## Acknowledgments

We thank the Western Regional Small Grains Genotyping Laboratory, specifically Travis Ruff and Marcus Hooker for the second Ion Torrent sequencing run, and to the staff and research personnel of the Central Research Institute for Field Crops (Ankara, Turkey) for their help.

## Supplementary Material

**Supplemental Figure 1.** Same as Figure 2 but inverted for susceptibility alleles. Enrichment dumbbell plot of enrichment of the susceptibility allele in landrace (red) and wild (blue) barley accessions. Percentage is included below each data point and difference between the two barley classes as well as which isolate the loci was identified with.

